# Brackish Beginnings: Rethinking the Role of Salinity in Shaping Mosquito Proboscis Morphology and Disease Risk

**DOI:** 10.64898/2026.01.28.702327

**Authors:** Charles Arokiyaraj, S Sreelakshmi, M Dharshini, Amit Kumar

## Abstract

Climate change driven saltwater intrusion is expanding coastal brackish water habitats, promoting the proliferation of salinity-tolerant mosquitoes such as *Culex sitiens*, a vector of lymphatic filariasis and Japanese encephalitis. This study investigated whether environmental salinity affects mosquito morphology, specifically proboscis length, a trait of ecological significance related to feeding efficiency and vector competence. Late fourth-instar larvae and pupae of *Cx. sitiens* were collected from habitats with varying salinity levels in the Muttukadu Backwater, Tamil Nadu, India, and reared under controlled conditions using habitat specific water. Adult female body and proboscis lengths were measured, and water quality parameters were analysed to characterize environmental variation. Statistical analyses (Welch’s ANOVA, regression, and ANCOVA) revealed a significant positive relationship between salinity and body length (R^2^ = 0.94, p=0.0003) as well as with proboscis length (R^2^ = 0.90, p=0.001). Additionally, ANCOVA indicated that the proboscis elongation remained significant after adjusting for body length (F□, □□□ = 32.36, p < 0.001, partial η^2^ = 0.257). This confirmed that the salinity exerts an independent effect on this morphological trait. These findings provide the first field-based evidence that the environmental salinity drives proboscis elongation in *Cx. sitiens*, indicating an adaptive response under saline stress. This may have implications for disease transmission in climate-affected coastal regions.

## 1. Introduction

Brackish water environments are ecologically significant transition zones that support biodiversity and shape the ecology of disease vectors such as mosquitoes (Bruland, 2008). In coastal regions, these habitats promote the emergence and persistence of salinity-tolerant mosquito species. Under dynamic conditions, including increased rainfall, tidal intrusion, and anthropogenic activities, some freshwater mosquito species can adapt to brackish environments (Balasubramanian et al., 2021). To cope with such abiotic stressors, mosquitoes may exhibit specific morphological adaptations, which remain largely underexplored.

Among the major drivers of this environmental shift, climate change-induced saltwater intrusion has emerged as a critical factor. Saltwater intrusion into coastal habitats is an increasingly pervasive global phenomenon (Prusty and Farooq, 2020). This process expands brackish water bodies, which serve as breeding habitats for various mosquito species, thereby increasing the risk of mosquito-borne disease transmission in vulnerable coastal populations (Arokiyaraj et al., 2025).

*Culex sitiens* Wiedemann, 1828, is a mosquito species widely distributed across tropical and subtropical regions, closely associated with brackish and saline water bodies (Rattanarithikul et al., 2005). It is a medically important vector known to transmit *Brugia malayi*, the causative agent of lymphatic filariasis, and Japanese encephalitis virus (JEV) (Bordoloi et al., 2021). *Culex sitiens*, a nocturnal species, exhibits high biting activity and thrives in diverse coastal habitats such as ponds, ditches, crab holes, salt marshes, and mangroves under a wide range of salinity conditions (Dash, 2011). Biting rates can reach up to 108 mosquitoes per person per hour in some regions, making it both a significant nuisance and a public health concern (Prummongkol et al., 2012). Globally, over 657 million people across 39 countries require preventive chemotherapy for lymphatic filariasis, and more than 3 billion are at risk of JEV infection, underscoring the vectorial importance of this species (WHO, 2024a, b). With ongoing climate change accelerating the expansion of inland saline habitats, the public health threat posed by this species is expected to increase.

Environmental factors such as salinity gradients, wind dynamics, and tidal fluctuations play a crucial role in shaping mosquito distribution, abundance, and phenotypic plasticity in coastal ecosystems (Zittra et al., 2017). These abiotic stressors can influence mosquito morphology, behaviour, and life-history traits, including development rate, body size, longevity, host-seeking, and feeding efficiency (Ramasamy and Surendran, 2012; Chandrasegaran et al., 2020). While the effects of temperature, humidity, and precipitation on mosquito traits have been widely studied, the influence of salinity on mosquito morphology remains poorly understood.

Among mosquito morphological traits, the proboscis has a direct influence on host penetration efficiency, feeding success, and consequently, pathogen transmission potential. Since these functions are critical under salinity-induced stress conditions, proboscis length was selected as a representative and ecologically meaningful trait for assessing salinity-driven morphological plasticity. Despite increasing recognition of saline adaptation, the morphological correlates of this process remain under characterized.

To address this knowledge gap, we investigated the morphological characters of *Cx. sitiens* developing under varying environmental salinity. We specifically focused on the proboscis length, a key trait linked to feeding efficiency and vector competence. We selected *Cx. sitiens* for two main reasons: (i) its medical importance as a vector and (ii) it demonstrated ability to tolerate a broad range of salinity conditions (Bordoloi et al., 2021; Jude et al., 2012). We hypothesized that increasing environmental salinity during larval development would induce elongation of the proboscis in female *Cx. sitiens* independent of changes in overall body size. This study provides new insights into salinity-driven phenotypic plasticity in a coastal mosquito species and contributes to a better understanding of local adaptation mechanisms critical for developing ecologically informed vector control strategies. Understanding these morphological adjustments provides insight into how mosquitoes may adapt to future salinization of coastal habitats.

## 2. Materials and Methods

### 2.1. Study area

As part of coastal mosquito monitoring program, brackish water mosquito surveillance was conducted across 30 locations along the southeast coast of Tamil Nadu, India. Based on this long-term dataset, two sites with consistently high *Cx. sitiens* abundance were identified as potential study locations and selected for field sampling. These sites were located within the Muttukadu Backwater (12.7959°N, 80.2461°E), a bar-built estuarine system in Covelong, Tamil Nadu, situated approximately 36 km south of Chennai. The estuary, covering an area of ∼3 km^2^, extends ∼15 km in a north–south direction with a width of 800–1050 m and a maximum central depth of 2 m, although the majority of the area remains shallow (≤1 m). It is seasonally reconnected to the Bay of Bengal through sandbar erosion caused by inland water inflow. The region supports multiple human activities, including fishing and boating. The two sampling sites within the estuary were designated as S1 (Kelambakkam) and S2 (Covelong), and are indicated in red on the study area map (Fig. 1).

**Fig. 1.**
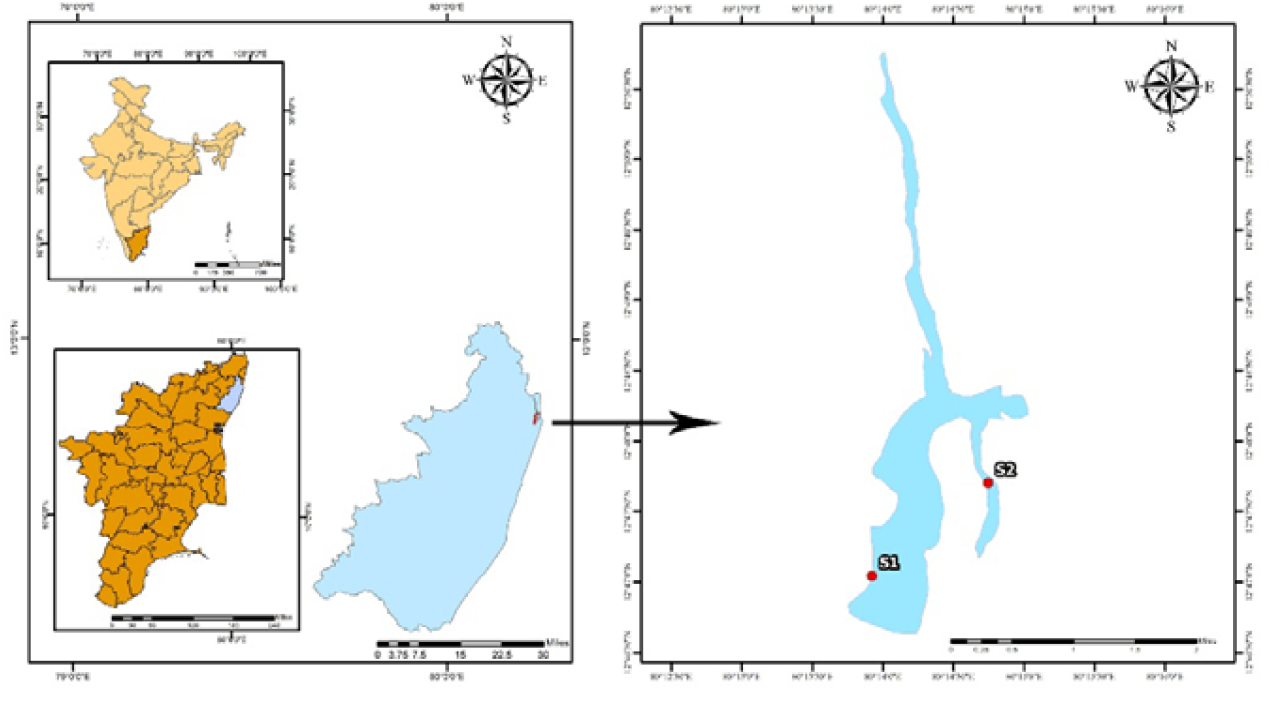
Location map of the study area. The inset maps show (top left) India with the state of Tamil Nadu highlighted, and (bottom left) Tamil Nadu with the Chengalpattu district highlighted in blue. The main map (right) displays the Muttukadu backwater system along the southeastern coast of India. Two sampling stations are marked: S1 – Kelambakkam and S2 – Covelong.

### 2.2. Mosquito collection

Mosquito sampling was conducted temporally from May 2024 to July 2025 at the study sites. Late fourth-instar larvae and pupae of *Cx. sitiens* were collected from these sites. These advanced developmental stages were chosen to minimise laboratory-induced effects and better capture the influence of natural environmental conditions. Collected specimens were transported in habitat water to the Vector-Borne Disease Research Facility at Centre for Climate Change Studies, Sathyabama Institute of Science and Technology, Chennai. In the laboratory, larvae and pupae were reared to adulthood in source water under controlled conditions: temperature 28□± □2□°C, relative humidity 75□± □2%, and a 12:12 h light-dark photoperiod. All specimens emerged as adults within 48 hours. Species identity was confirmed by examining key meristic and morphometric traits as described in Tyagi et al. (2025). In addition, selected specimens were validated using molecular identification targeting the mitochondrial cytochrome c oxidase subunit I (COI) gene following Folmer et al. (1994). The resulting sequences were deposited in GenBank (Accession Nos. PV946892 and PV946893).

### 2.3. Environmental parameter assessment

*In situ* water quality parameters, including salinity, total dissolved solids (TDS), pH, and temperature, were measured during specimen collection at each site using a waterproof multiparameter handheld instrument (Lovibond®, Germany). Additional water samples were also collected in sterile containers to measure inorganic phosphate, nitrite, and nitrate using spectrophotometric methods, following the standard procedures described by Strickland and Parsons (1972).

### 2.4. Body and proboscis morphometry

Both body and proboscis lengths of adult female mosquitoes were measured using a stereo zoom microscope (SMZ25-Nikon, Japan) equipped with NIS-Elements imaging software. To evaluate the influence of salinity, specimens were grouped into four categories based on habitat salinity: low (0–10 ppt), mild (11–20 ppt), moderate (21–30 ppt), and high (>30 ppt). For this study, representative salinity levels included 3 and 6 ppt (low), 13 ppt (mild), 21 and 29 ppt (moderate), and 31 and 34 ppt (high). Each salinity group included a pooled sample collected from study sites at different periods. Mosquitoes were anaesthetised with CO□ and mounted laterally on glass slides to ensure consistent imaging. Proboscis length was measured from its base (point of attachment to the head) to the tip, while body length was measured from the anterior tip of the head to the posterior end of the abdomen. Both proboscis and body lengths were recorded for each adult female mosquito to distinguish the specific effect of salinity on proboscis morphology from general size variation. All measurements were recorded in micrometres (Fig. 2a). Triplicate measurements were taken for each individual to ensure accuracy and reproducibility.

**Fig. 2.**
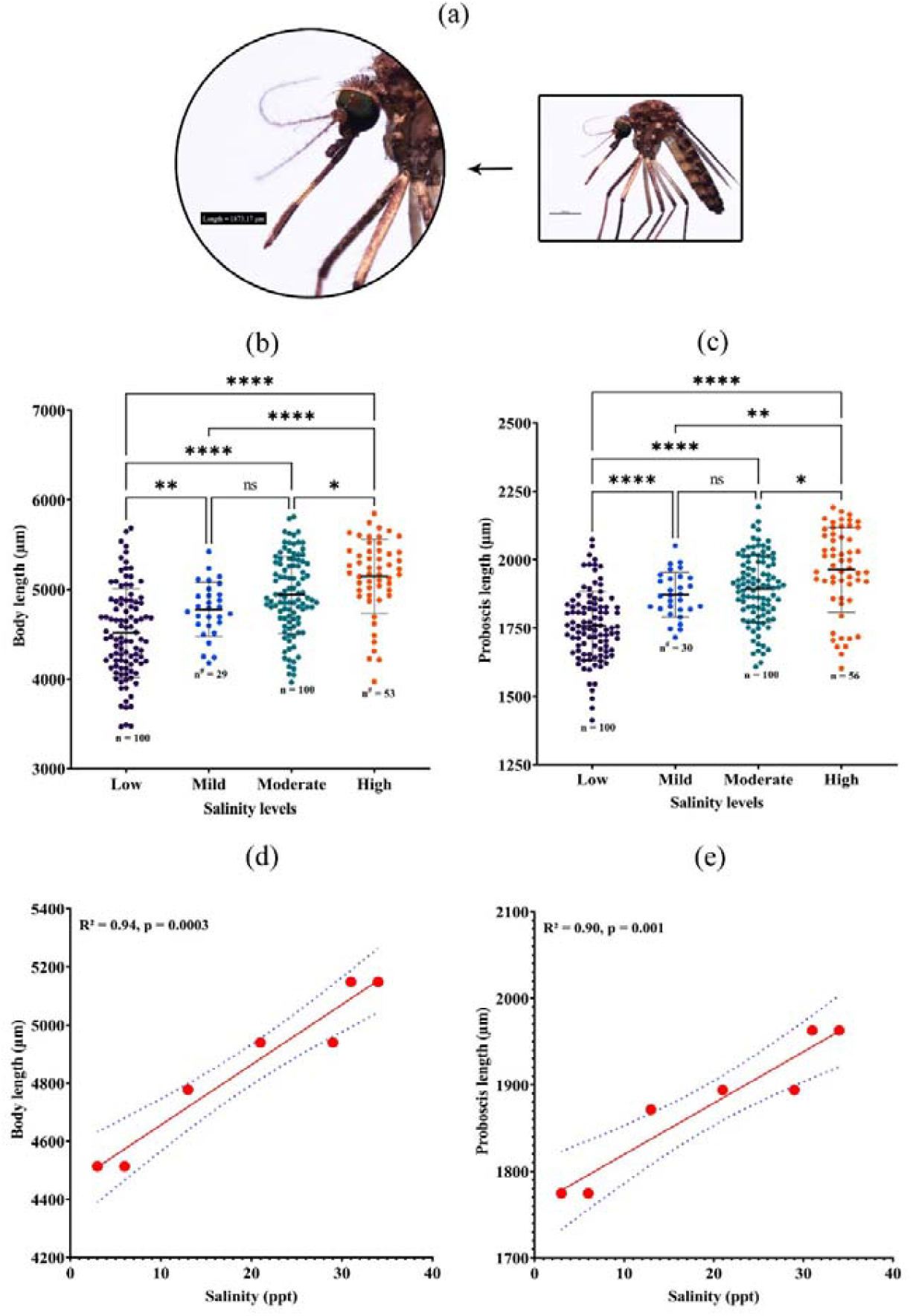
Workflow and salinity-associated changes in body and proboscis lengths in *Cx. sitiens*. (a) Adult female *Cx. sitiens* imaged under a stereomicroscope, showing the elongated body and proboscis lengths. The inset illustrates the measurement process of mosquito structures (e.g., proboscis) using calibrated imaging software. (b) and (c) Scatter dot plots illustrating the variation in body length and proboscis length (µm) of female *Cx. sitiens* across four salinity levels (Low, Mild, Moderate, High). Horizontal bars represent mean ± SD, and sample sizes (n) are indicated below each group. ‘#’ indicates outliers excluded from analysis within each group (ranging from 1 to 3 individuals). Statistical significance among groups was assessed using Welch’s ANOVA. Asterisks indicate significance levels: p < 0.05 (*), p < 0.01 (**), p < 0.001 (***), p < 0.0001 (****); ns = not significant. (d) and (e) Linear regressions showing positive associations between salinity (ppt) and both body length (R^2^ = 0.94, p = 0.0003) and proboscis length (R^2^ = 0.90, p = 0.001), indicating a progressive elongation of morphological traits with increasing salinity levels. Red dots represent group means; solid red lines show fitted regression models, and blue dashed lines denote 95% confidence intervals.

### 2.5. Statistical analysis

To evaluate whether salinity influenced proboscis length independently of overall body size, a series of statistical tests were conducted. Initially, Welch’s ANOVA with Dunnett’s T3 multiple comparison test was used to compare mean body and proboscis lengths among the four salinity groups, accounting for unequal sample sizes. Statistical significance was set at p <0.05. To explore relationships between individual abiotic factors and mosquito morphometrics, simple linear regression and Pearson correlation analyses were performed, with body and proboscis lengths as dependent variables and abiotic parameters as independent predictors. Each model produced slope coefficients, R^2^ values, and p-values to assess the strength and significance of associations. To determine whether proboscis elongation persisted after adjusting for overall body size, an Analysis of Covariance (ANCOVA) was performed in IBM SPSS Statistics 26. In this model, salinity group was treated as a fixed factor, proboscis length as the dependent variable, and body length as a covariate. Before analysis, assumptions of normality and linearity were confirmed from residual plots. Levene’s test of Equality of Error Variances was violated (p = 0.021); however, given the reasonably balanced sample sizes, this model was considered robust to this deviation. Partial eta squared (η^2^) values were used to estimate effect sizes. Bonferroni-adjusted post-hoc pairwise comparisons were conducted wherever appropriate. The analyses and visualisations were carried out in GraphPad Prism version 9 for Windows.

## 3. Results

Body and proboscis lengths of *Cx. sitiens* females varied significantly across the salinity gradient. Mosquitoes emerging from high-salinity habitats exhibited the greatest body and proboscis lengths, whereas individuals from low-salinity environments showed the smallest values for both traits (Fig. 2b–c). Pairwise comparisons revealed highly significant differences between the low and moderate salinity groups (p < 0.0001) and between the low and high salinity groups (p < 0.0001). Significant differences were also detected between the moderate and high salinity groups (p < 0.05). In comparisons between the mild and high salinity groups, body length showed a highly significant increase (p < 0.0001), whereas proboscis length exhibited a significant but comparatively weaker difference (p < 0.01). No statistically significant difference was observed between the mild and moderate salinity groups.

Regression analyses demonstrated strong positive relationships between environmental salinity and both body length (R^2^ = 0.94, p = 0.0003) and proboscis length (R^2^ = 0.90, p = 0.001), indicating that salinity alone accounted for more than 90% of the observed variation in these morphological traits (Fig. 2c–d). In addition to salinity, total dissolved solids (TDS) and pH were also positively associated with body and proboscis lengths (Tables 1 and 2).

**Table 1.**
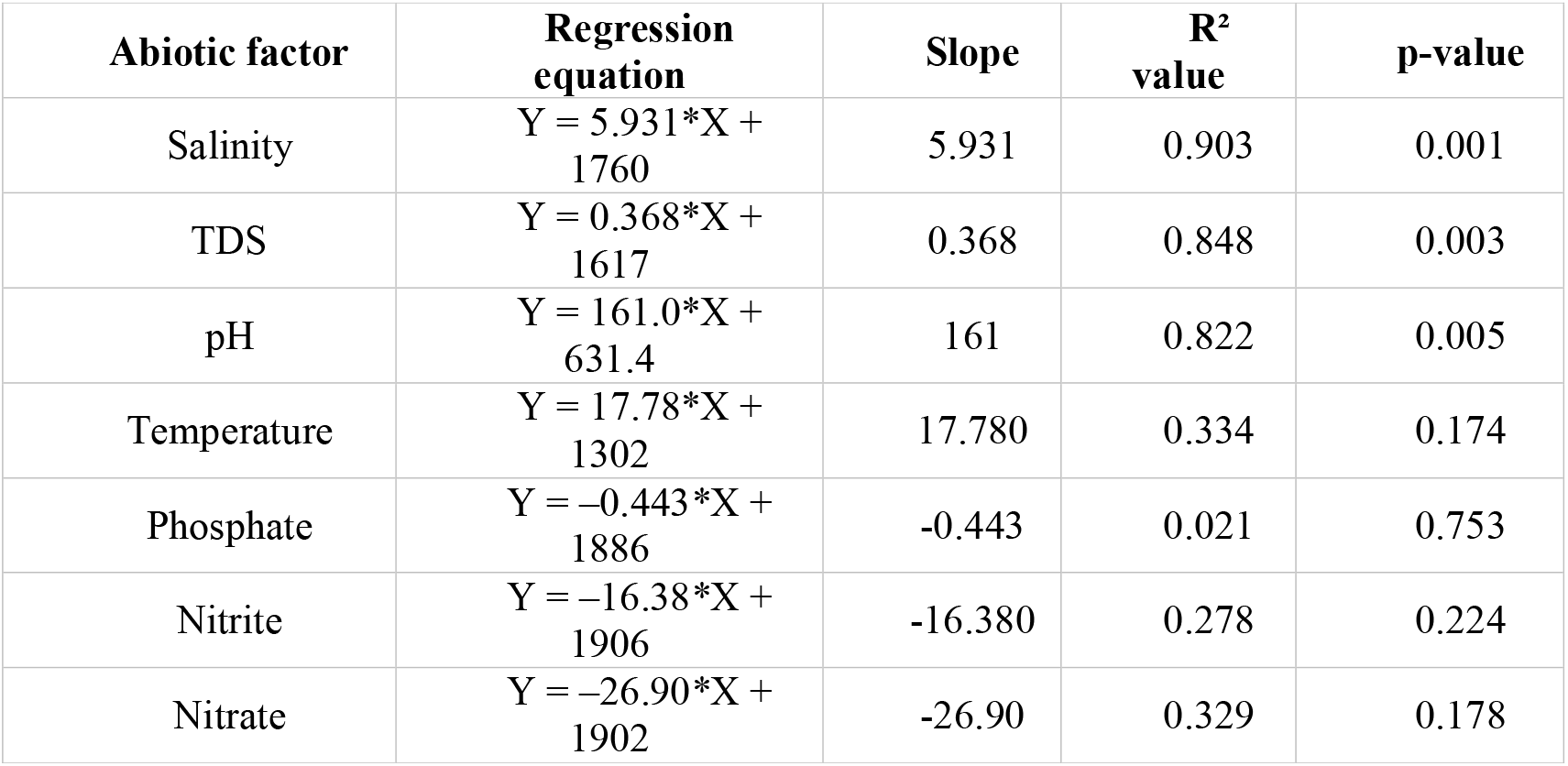
Simple linear regression analysis of abiotic factors influencing proboscis length in *Cx. sitiens*.

**Table 2.** Simple linear regression analysis of abiotic factors influencing body length in *Cx. sitiens*.

| Abiotic factor | Regression equation | Slope | R<sup>2</sup> value | p-value |
| --- | --- | --- | --- | --- |
| Salinity | $Y = 20.72 * X + 4449$ | 20.72 | 0.940 | 0.0003 |
| TDS | $Y = 1.236 * X + 3983$ | 1.236 | 0.817 | 0.0052 |
| pH | $Y = 549.0 * X + 609.8$ | 549.0 | 0.815 | 0.0053 |
| Temperature | $Y = 63.19 * X + 2812$ | 63.19 | 0.360 | 0.1546 |
| Phosphate | $Y = -1.396 * X + 4885$ | -1.396 | 0.018 | 0.7721 |
| Nitrite | $Y = -52.29 * X + 4950$ | -52.29 | 0.242 | 0.2622 |
| Nitrate | $Y = -88.43 * X +$ | -88.43 | 0.303 | 0.2004 |
|  | 4939 |  |  |  |

In contrast, water temperature, phosphate, nitrite, and nitrate concentrations showed no significant relationships with either body or proboscis length.

To assess whether changes in proboscis length occurred independently of overall body size, an analysis of covariance (ANCOVA) was performed with body length included as a covariate. This analysis revealed that proboscis length differed significantly among salinity groups even after adjusting for body length (F□, □□□ = 32.36, p < 0.001), with a large effect size (partial η^2^ = 0.257). These results indicate that proboscis elongation in *Cx. sitiens* from high-salinity environments cannot be explained solely by increases in overall body size and instead represents a distinct morphological response associated with saline habitat conditions (Table 3; Fig. 3).

**Table 3.**
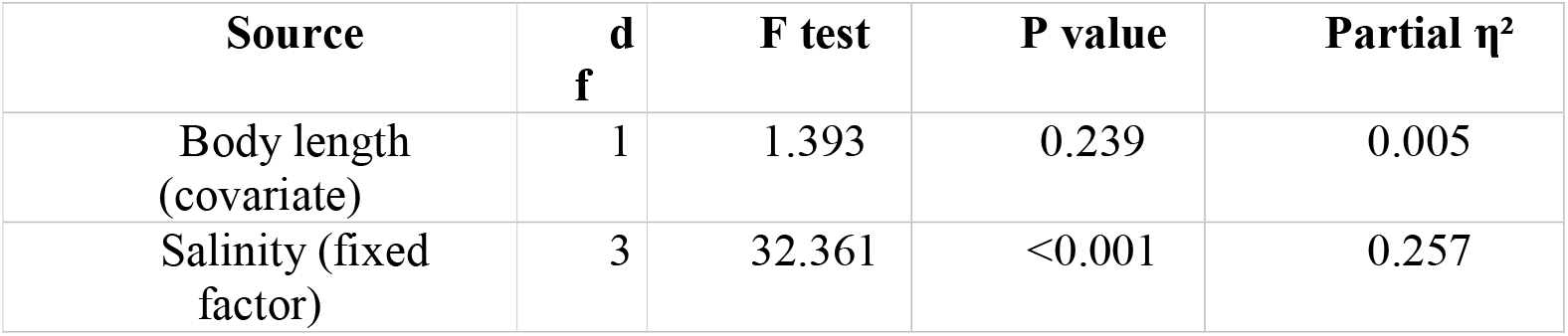
Results of ANCOVA testing the effect of salinity on proboscis length after adjusting for body length in *Cx. sitiens*.

**Fig. 3.**
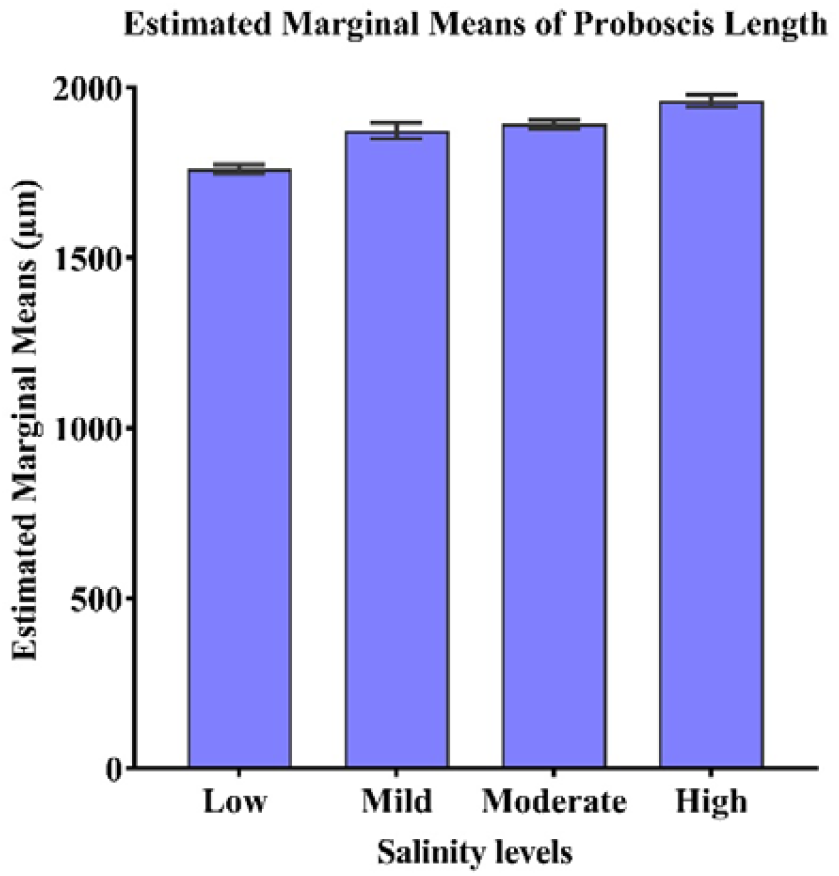
Estimated marginal means of proboscis length (µm) across four salinity groups after adjusting for body length using analysis of covariance (ANCOVA). Bars represent adjusted mean proboscis length ± SE, with covariates evaluated at the mean body length (4,743.89 µm). The increasing pattern of adjusted marginal means across salinity categories visually indicates a positive association between salinity and proboscis length, which was confirmed by ANCOVA as a significant main effect of salinity (F□3,281□ = 32.36, p < 0.001, partial η^2^ = 0.257).

## 4 Discussion

Our findings demonstrate that environmental salinity induces elongation of the proboscis in *Cx. sitiens* independent of body size, representing a novel field-based example of salinity-driven morphological plasticity in mosquitoes. Although proboscis morphology has been extensively investigated in lepidopterans to understand feeding adaptations (Vajna et al., 2021), its ecological and functional significance in mosquitoes remains largely underexplored. In mosquitoes, proboscis length is linked not only to feeding efficiency and vector competence but also serves as a useful taxonomic character and a potential indicator of physiological and developmental responses to environmental stressors.

Previous experimental studies have shown that temperature can influence proboscis length in mosquitoes (Agyekum et al., 2022). However, the influence of salinity as a developmental factor has received almost no attention. To our knowledge, this is the first field-based observation examining the relationship between environmental salinity and proboscis morphology in mosquitoes. Individuals emerging from high-salinity habitats exhibited significantly longer proboscises than those from low-salinity water. This pattern suggests that elevated salinity during larval development may induce proboscis elongation as an adaptive plastic response that enhances feeding efficiencies, and potentially survival under saline conditions. Because proboscis morphology can influence biting efficiency and host penetration, such salinity-induced morphological variation can have downstream effects on pathogen transmission potential.

These interpretations are supported by a strong positive correlation between salinity and proboscis length (R^2^ = 0.90, p = 0.001), with additional associations observed for TDS and pH, indicating that ionic composition collectively influences morphology. Other abiotic variables, such as temperature, phosphate, nitrite, and nitrate, showed no significant relationships. The observed elongation may therefore represent a functional adaptation that optimizes feeding efficiency and host penetration in saline environments, thereby enhancing vectorial capacity (Chandrasegaran et al., 2025).

Mechanistically, the plasticity observed in *Cx. sitiens* is likely mediated by osmoregulatory stress experienced during larval development in high-salinity environments. Such stress is known to influence hormonal pathways in insects, particularly the ecdysone signaling cascade that regulates growth and morphogenesis. Although speculative, this pattern could be mediated by ecdysone-regulated developmental adjustments under osmotic stress. Ecdysone-mediated modulation of developmental timing and resource allocation remains a plausible mechanism underlying salinity-driven proboscis elongation. This reflects a broader physiological principle wherein stress-induced endocrine adjustments facilitate ecological plasticity and enhance adaptation to challenging habitats (Hun et al., 2022).

These findings highlight that salinity-induced morphological plasticity may affect the feeding success and pathogen transmission potential of coastal mosquitoes. With global sea-level rise and anthropogenic alteration, understanding such adaptive responses is critical for predicting shifts in mosquito distribution, host interactions, and vectorial capacity. Integrating salinity dynamics into mosquito surveillance and vector control strategies may therefore be essential for mitigating disease risks in climate-vulnerable coastal zones.

Wing morphology and other flight-related traits are known to respond to environmental gradients across wider geographic ranges. In this study, we focused solely on proboscis within a single coastal habitat. Earlier studies have reported that ecological factors can influence mosquito morphometric traits, such as wing pattern, cuticle structure, body size, and proboscis length. However, the underlying developmental mechanisms remain poorly understood (Agyekum et al., 2022; Barr et al., 2023; Laojun et al., 2025; Sivabalakrishnan et al., 2025). Environmental stressors can trigger plastic responses across mosquito taxa, though the magnitude and nature of these responses are likely species-specific. Our findings align with the principles of ecological developmental biology, which propose that environmental cues modulate phenotypic outcomes during growth (Sivabalakrishnan et al., 2023; Abdulloh et al., 2024). Although correlation does not prove causation, the consistency of our results suggests that salinity plays a meaningful role in shaping mosquito morphology in coastal ecosystems.

Future studies should investigate the physiological and molecular mechanisms underlying salinity-mediated morphological changes. These insights can refine ecological models of vector adaptation and resilience. Integrating these mechanistic insights with environmental monitoring may improve predictions of mosquito population dynamics and disease transmission under changing coastal conditions. Understanding salinity-mediated morphological plasticity thus offers an early warning framework for anticipating vector behavioural shifts and optimizing control strategies under accelerating coastal salinization.

## Acknowledgements

The authors are grateful to the management of the Sathyabama Institute of Science and Technology for providing the necessary facilities to conduct this research work. The authors are thankful to the Indian Council of Medical Research (ICMR) for providing research funds to support this work (No. 6/9-7(328)/2023/ECD-II).

